# Representative vs. Load-bearing Layers: A Dissociation in Genomic Foundation Models

**DOI:** 10.64898/2026.07.16.739040

**Authors:** Yoonjin Cho, Min Seok Kim, Sangwoo Kim

**Affiliations:** Yonsei University College of Medicine, Seoul, Republic of Korea; Department of Biomedical Systems Informatics, Yonsei University College of Medicine, Seoul, Republic of Korea

## Abstract

Downstream use of genomic foundation models follows one of three conventions: aggregating representations across all layers (Pearce et al., 2026), defaulting to the last hidden state as a fixed feature extractor (Dalla-Torre et al., 2024), or picking a single intermediate layer via mechanistic-interpretability tooling (Brixi et al., 2026). None of these examines which layer a joint classifier actually relies on. We probe this question with a minimal training-free scalar, ∥Δ*h_ℓ_*∥_2_, defined as the L2 norm of the per-layer hidden-state shift at the variant token. We evaluate it on 8,008 ClinVar (Landrum et al., 2018) single-nucleotide variants in NT-v2 500M (Dalla-Torre et al., 2024), a masked language model (MLM), and Evo 2 7B (Brixi et al., 2026), a causal language model (CLM) with a hyena/attention hybrid. In both models the layer with peak single-feature AUROC (the *representative* layer) is not the layer a joint multi-layer classifier most depends on (the *load-bearing* layer, identified by leave-one-layer-out ablation drop and concordant with |SHAP| (Lundberg & Lee, 2017)). Representative layers sit mid-network in both models, whereas load-bearing depth lies at opposite ends of the depth axis: mid-shallow in the MLM, and deep in the CLM hybrid. The dissociation has direct downstream consequences. In NT-v2, a 1-dimensional mid-layer scalar exceeds the canonical 1024-dimensional last-layer mean-pool baseline by +0.049 AUROC. In Evo 2, the 4096-dimensional mean-pool is competitive with the joint ∥Δ*h_ℓ_*∥_2_ feature, so standard last-layer pooling leaves variant-relevant signal untapped specifically in MLM-based pipelines.

## 1. Introduction

Layer-wise use of genomic foundation models (FMs) has so far split across three conventions. Aggregation methods pool features across all layers (Pearce et al., 2026), treating layer identity as a nuisance dimension; mechanistic analyses pick a single intermediate layer based on the density of biologically meaningful features (Brixi et al., 2026; Templeton et al., 2024); and NT-v2–style downstream featureextraction pipelines default to the last hidden state (Dalla-Torre et al., 2024). None of these conventions assesses the question that separates them: *which layer does the joint model actually rely on, and is it the same layer a univariate probe identifies as carrying the strongest signal?*

In this paper, we demonstrate that this alignment does not hold across the two genomic FMs studied here. The layer that maximises single-feature variant discriminability is *not* the layer the joint multi-layer classifier most depends on, in both NT-v2 (a masked language model, MLM) and Evo 2 (a causal language model, CLM, with a hyena/attention hybrid backbone). We diagnose this with a minimal training-free probe and two operational metrics introduced in §2: per-layer single-feature AUROC, which we call the *representative* signal (“where does the signal live?”), and leave-one-layer-out ablation drop on a joint multi-layer classifier, which we call the *load-bearing* signal (“what does the joint model rely on?”), with |SHAP| (Lundberg & Lee, 2017) as a concordant attribution.

### Contributions

We demonstrate a representative–load-bearing dissociation in two genomic FMs (§3–§4): the layer with the strongest univariate signal is not the layer the joint classifier most relies on. We further observe a cross-architectural depth shift in which relative load-bearing depth is mid-shallow under the MLM and deep under the CLM hybrid, while representative depth remains mid-network in both architectures; because objective, capacity, and architecture covary across the two models, we treat this shift as observational rather than causal. The dissociation has a downstream consequence specific to MLMs: in NT-v2 a 1-dimensional mid-layer scalar exceeds the canonical 1024-dimensional last-layer mean-pool by +0.049 AUROC at zero added compute, whereas in Evo 2 the mean-pool remains comparable to the joint ∥Δ*h*_*ℓ*_∥_2_ feature (App. F). This asymmetry reflects MLM-specific representation structure rather than a generic genomic-FM property.

Our scope is intentionally narrow: one task (variant pathogenicity), two main models with one capacity-limited control, and linear readouts. Within that scope, the within-model dissociation replicates across four scalar reductions of Δ*h*_*ℓ*_ (App. H).

### Related work

Layer-wise probing has a long tradition in NLP: the logit lens (nostalgebraist, 2020) and TunedLens (Belrose et al., 2023) read intermediate hidden states through the language-model head, Tenney et al. (2019) dissect BERT depth into a classical NLP pipeline, and recent evidence indicates that last-layer features are systematically suboptimal for downstream tasks (Skean et al., 2025). In genomic FMs, prior work either pools across layers (Pearce et al., 2026) or selects single intermediate layers via mechanistic-interpretability tooling (Brixi et al., 2026; Templeton et al., 2024). Our contribution is to formalise these as two distinct questions — which layer is most *decodable* versus which layer is most *relied upon* — and to show that they identify operationally distinct layers in the genomic FMs we study.

## 2. Probe and setup

For each variant we run two forward passes through a frozen genomic foundation model, one on the reference sequence and one on the alternate (Fig. 1). With 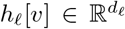 the hidden state at layer *ℓ* and the variant-token position *v*, the probe is

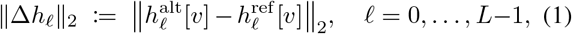

a minimal, training-free, rotation-invariant, head-decoupled scalar. Three alternative reductions (cosine, L1, directional projection) yield the same within-model dissociation (App. H); construction details are in App. A.

**Figure 1.**
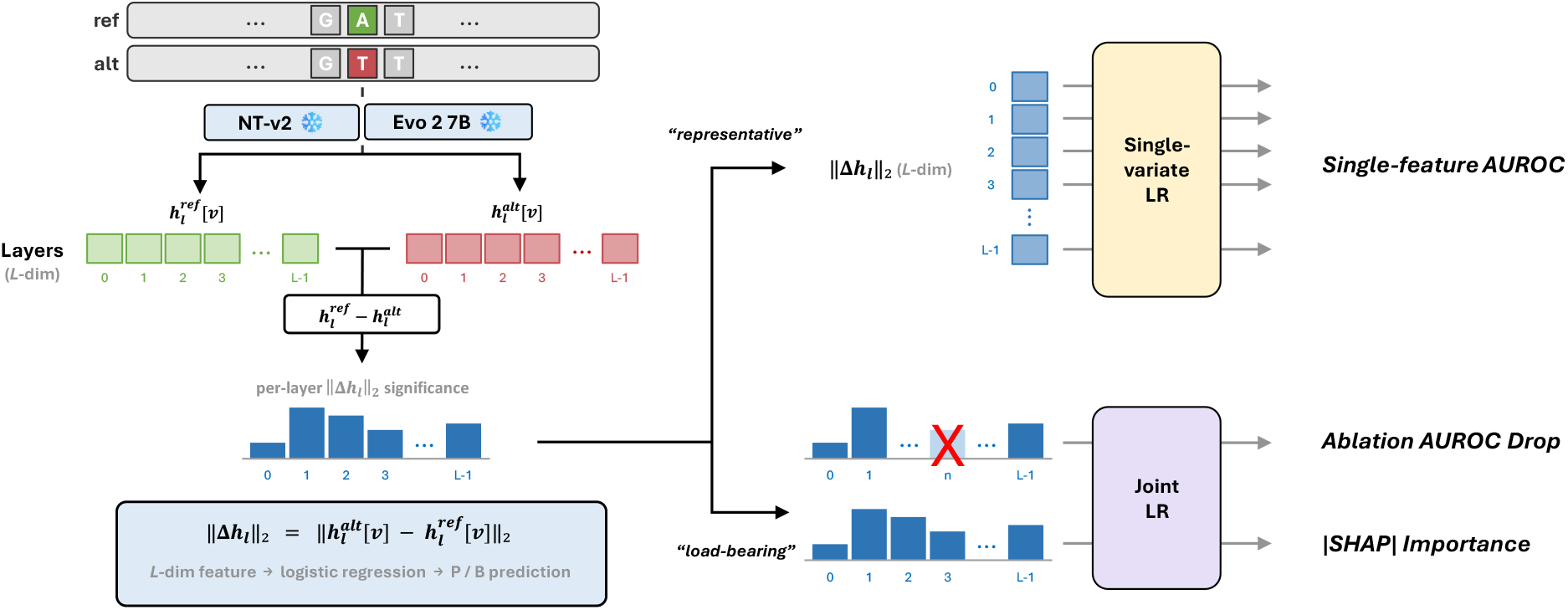
Method overview and the two metric axes. **Left:** matched ref/alt forward passes through a frozen genomic FM (NT-v2 or Evo 2). At the variant-token position *v* we compute ∥Δ*h*_*ℓ*_∥_2_, the L2 norm of the per-layer hidden-state difference; stacking across layers gives an *L*-dim feature (*L*=30 for NT-v2, *L*=32 for Evo 2; index *ℓ*=0, …, *L* − 1). **Right:** the two operational axes. The *representative* axis (top) reports per-layer single-feature AUROC; the *load-bearing* axis (bottom) reports leave-one-layer-out ablation drop (X marks the dropped layer) and |SHAP| on the joint classifier (§3–§4). The two axes need not select the same layer.

### Two operational definitions

The **representative** layer is the per-layer single-feature AUROC peak (*“where does the signal live?”*). The **load-bearing** layer is the layer with the largest leave-one-layer-out ablation drop on the joint multi-layer classifier, concordant with |SHAP| (Lundberg & Lee, 2017) (*“what does the joint model rely on?”*). A single layer need not answer both questions, and we show in §3 and §4 that the two definitions identify operationally distinct layers in both architectures.

We use 8,008 ClinVar (Landrum et al., 2018) single-nucleotide variants across 15 high-penetrance cancer-associated genes (Pathogenic / Likely Pathogenic, P/LP, *n*=3,514; Benign / Likely Benign, B/LB, *n*=4,494; App. D). Models are summarised in Table 1: **NT-v2 500M** (Dalla-Torre et al., 2024) and **Evo 2 7B** (Brixi et al., 2026) are the two main models; **HyenaDNA-large** (Nguyen et al., 2023) is a smaller capacity-limited control based on Hyena operators rather than attention, reported separately in App. K. Throughout, *ℓ* indexes post-block hidden states; NT-v2 includes the input embedding as *ℓ*=0 (indices *ℓ*=0, …, 29) while Evo 2 follows its paper’s convention over the 32 post-block outputs (*ℓ*=0, …, 31); full details in App. A. All models are frozen.

**Table 1.** Models evaluated. *L* is the number of hidden-state layers we probe under each model’s own indexing convention (App. A).

| Model | Params | Objective | Backbone | Tokens | $L$ |
| --- | --- | --- | --- | --- | --- |
| NT-v2 500M | 500M | MLM | Transformer | 6-mer | 30 |
| Evo 2 7B | 7B | CLM | Hyena/attn hybrid | single-base | 32 |
| HyenaDNA-large | 6.6M | CLM | Hyena | single-base | 10 |

We use the same variant split and preprocessing for every layer and architecture, so differences across layers reflect representation depth rather than changes in the downstream protocol. Per-layer and joint AUROCs use stratified 10-fold logistic regression with no tuning; bootstrap CIs, DeLong tests, and three confound controls (substitution-stratified AUROC, within-gene label permutation, gene-balanced sub-sampling) all pass (App. C–L).

## 3. Identifying representative layers

To locate where the variant signal is most visible, we first evaluate each layer independently, before asking how layers behave when combined in a joint classifier. This per-layer view gives the simplest operational definition of a representative layer: the depth at which the ref/alt perturbation is most decodable on its own. In both models we observe a broad mid-network band where this scalar separates pathogenic from benign variants, but the two architectures behave differently near the output. NT-v2 loses the signal near the final layer, whereas Evo 2 retains a usable last-layer readout. This contrast motivates the later distinction between representative layers and joint-model reliance.

### Per-layer AUROC profile

For each layer we fit a stratified 10-fold logistic regression on the single feature ∥Δ*h*_*ℓ*_∥_2_ and report out-of-fold AUROC. The profile (Fig. 2) is consistent across architectures: a sharp rise out of the input embedding, a broad mid-network plateau, and decay or collapse at the final layer. NT-v2 plateaus over L9–L26 (≥ 0.90), peaks at ∥Δ*h*_15_∥_2_=0.930, and collapses near chance level at the final block (∥Δ*h*_29_ ∥_2_=0.550; 5 of 14 genes below chance under leave-one-gene-out; App. D). Evo 2 peaks at ∥Δ*h*_8_∥_2_=0.855 and decays more gently to ∥Δ*h*_31_∥_2_=0.750 (Δ=+0.105, no chance-level collapse). We interpret the NT-v2 collapse (§5) as a combination of final-layer basis rotation and MLM commitment to the reconstruction subspace. Both peak-vs-last gaps are highly significant (*p<*10^−50^ paired DeLong; 100% of paired bootstrap resamples; App. C).

**Figure 2.**
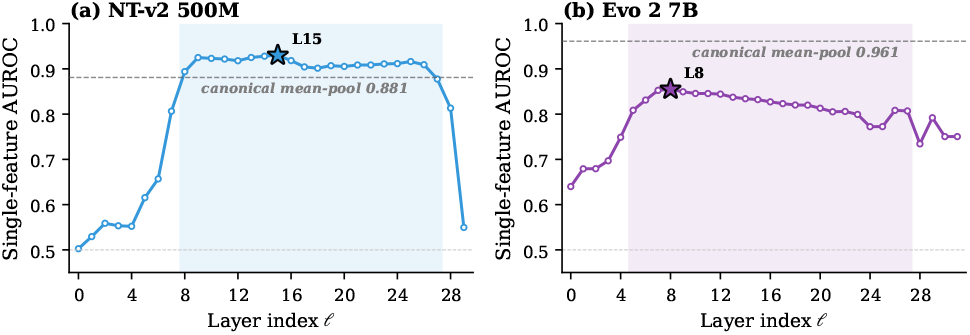
Per-layer single-feature AUROC. Out-of-fold AUROC of a 10-fold logistic regression on the single feature ∥Δ*h*_*ℓ*_∥_2_. The filled star marks the representative layer; the grey dashed line marks the canonical last-layer mean-pool baseline; the lower grey dotted line marks chance. **(a)** NT-v2 500M: peak L15 (0.930) exceeds mean-pool (0.881). **(b)** Evo 2 7B: peak L8 (0.855) sits below mean-pool (0.961).

### Comparison with the canonical last-layer mean-pool baseline

As a common feature-extraction baseline in NT-v2–style downstream benchmarks (Dalla-Torre et al., 2024), the last hidden state is mean-pooled over the input window and fed to a supervised head. Evo 2’s own downstream is instead likelihood scoring, but for a like-for-like comparison of layer-wise readouts we apply the same mean-pool protocol to both architectures on the same split (full Evo 2 numbers in App. F, Tab. A3); the two models behave qualitatively differently:

### NT-v2 (MLM)

The canonical 1024-dimensional meanpool last-layer baseline reaches 0.881 AUROC (Fig. 2a, grey dashed). The 1-dimensional mid-layer scalar ∥Δ*h*_15_∥_2_=0.930 exceeds this 1024-dimensional protocol by +**0.049** at a 1024× feature-dimensionality reduction.

### Evo 2 (CLM hybrid)

The canonical 4096-dimensional mean-pool last-layer baseline reaches 0.961 AUROC, comparable to (and slightly above) the joint 32-dimensional ∥Δ*h*_*ℓ*_∥_2_ feature (0.926). For Evo 2 the canonical meanpool readout is therefore not impaired; the corresponding fine-tune-free single-scalar comparison ( ∥Δ*h*_8_∥_2_=0.855 vs. ∥Δ*h*_31_∥_2_=0.750, Δ=+0.105) shows the within-architecture peak-vs-last gap, not a mean-pool deficiency.

### Architectural asymmetry

The +0.049 win is MLM-specific. NT-v2’s final-layer scalar collapses near chance, and the appendix strengthens the interpretation that this is not a single-gene artifact but a final-layer basis-rotation / reconstruction-subspace effect. The NT-v2 final layer shows a small sign-flipped residual in the univariate test (App. B), falls below chance in 5 of 14 evaluable genes under LOGO (App. D), and degrades the canonical mean-pool readout relative to the 1-dimensional mid-layer scalar (App. E). This is the pattern expected if MLM training rotates or compresses variant-relevant variance into directions poorly aligned with the final reconstruction readout (App. J). Evo 2 does not show this collapse: its final layer remains task-informative, so the canonical mean-pool stays competitive (App. F). Stacking ∥Δ*h*_*ℓ*_∥_2_ across all *L* layers raises the joint AUROC to 0.962/0.926 (NT-v2 / Evo 2), within ∼ 0.04 of the cross-layer covariance aggregator EVEE (Pearce et al., 2026) (0.997 on a different split) at the cost of one scalar per layer. This still gives a representative-layer view of where the signal lives, but not yet of which layer the joint model relies on.

## 4. Identifying load-bearing layers

When the joint multi-layer ∥Δ*h*_*ℓ*_∥_2_ feature (30-dimensional for NT-v2, 32-dimensional for Evo 2) is fit by logistic regression, which layer does the classifier most rely on? We identify the load-bearing layer by the largest leave-one-layer-out AUROC drop after refitting on the remaining *L* − 1 layers, with LightGBM SHAP as a concordant attribution (§2; App. A).

For NT-v2 the load-bearing layer is L9: both metrics rank L9 first (ablation drop +0.0027, |SHAP| 2.81, joint LR coefficient +4.21). The representative L15, despite the strongest single-feature AUROC (0.930), drops joint AUROC by only +0.0001 when removed. For Evo 2 the load-bearing layer is L29 (ablation +0.0069, ∼3× the runner-up); |SHAP| peaks nearby at L27, placing both attributions in the same last-five-layer band, far from representative L8 (Fig. 3b). Across layers, the attribution profiles are positively correlated (Spearman 0.78 NT-v2, 0.65 Evo 2).

**Figure 3.**
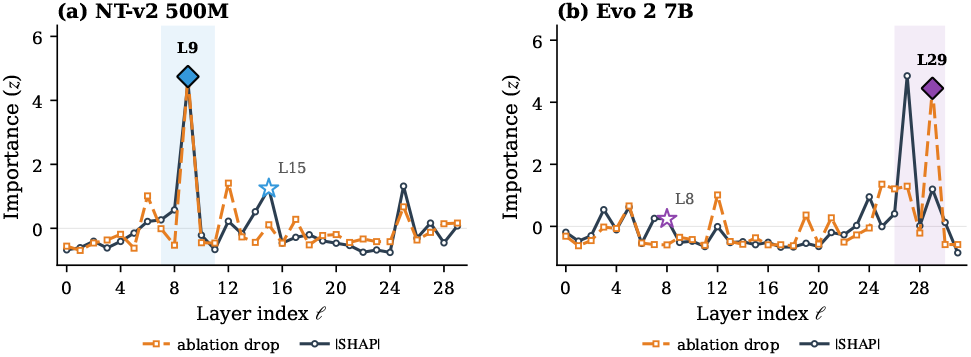
Per-layer joint-classifier importance. Tree-SHAP mean |SHAP| (navy solid) and leave-one-layer-out ablation drop (orange dashed), *z*-scored within architecture. The open star marks the representative layer; the filled diamond marks the load-bearing layer; the pink band highlights the load-bearing region. **(a)** NT-v2: load-bearing L9 vs. representative L15. **(b)** Evo 2: load-bearing L29 vs. representative L8.

### Suppressor layers

Layerwise contribution direction also includes a minor suppressive pattern in the joint LR coefficients. In this pattern, a layer can have high single-feature AUROC yet receive a negative coefficient once neighbouring layers are present, meaning that the joint model uses it to subtract or counterbalance variance that is otherwise predictive in isolation. This is consistent with the joint classifier recruiting these layers as a generic-perturbation-magnitude subtractor rather than as a variant-specific encoder: the layer carries a mixture of variant signal and a substitution-bias confound that scales with any sequence change, and the joint model removes the latter by entering the layer with a negative coefficient against positive load-bearing contributions. We treat this as a secondary effect rather than the main load-bearing result, with detailed discussion in App. I.

### Cross-architectural pattern

To compare models with different numbers of layers, we report depth as a relative position in the stack, from 0 at the input side to 1 at the final block. Representative depth is mid-network in both architectures (NT-v2 L15 at relative depth 0.52; Evo 2 L8 at 0.25), so the layer that is most decodable on its own sits in a broadly similar part of the computation despite the architectural differences. Load-bearing depth instead separates strongly in raw layer index and, more importantly, occupies opposite ends of the relative-depth axis: NT-v2 relies on a mid-shallow MLM layer (L9 at 0.31, before the representative layer), whereas Evo 2 relies on a deep CLM-hybrid layer (L29 at 0.91, near the final block). This contrast is the main structural result of the paper: the most visible variant signal is not enough to determine which layer the full classifier depends on once all layers are available together. The roughly twenty-layer raw-index gap should therefore be read through the normalized-depth comparison, where the load-bearing layer moves from the earlier third of the MLM stack to the final tenth of the CLM-hybrid stack. The within-model dissociation holds in both cases, while the cross-architectural direction is observational because objective, capacity, sequence mixing, and data covary (§5).

## 5. Discussion

### Mechanism of the dissociation

A layer is *representative* when its signal alone discriminates the task, and *loadbearing* when removing it from the joint feature drops classifier AUROC the most. The representative layers (Evo 2 L8, NT-v2 L15) are strong individually yet dispensable under leave-one-layer-out ablation once the joint classifier is refit on the remaining layers. The load-bearing layers (Evo 2 L29, NT-v2 L9) carry contributions that the joint classifier weights heavily and that ablation cannot replace; this is the operational signature of *reliance*. Standard single-layer probing protocols implicitly conflate these two roles because they treat the most decodable layer as the layer that matters functionally. Distinguishing them requires an analysis that keeps the joint classifier in the loop, such as leave-one-layer-out ablation together with attribution across the full layer stack.

### Cross-architectural depth allocation

Representative depths are mid-network in both architectures (relative depths 0.52 NT-v2, 0.25 Evo 2), whereas load-bearing depths lie at opposite ends of the normalized layer axis: NT-v2 relies on an early layer, about one-third through the stack (0.31), while Evo 2 relies on a late layer, near the output end (0.91). A capacity-limited control model (HyenaDNA-large, ∼6.6M params) collapses load-bearing to the input embedding (Table 2; App. K). Thus the dissociation is consistent across all three points, but load-bearing *depth* appears shaped jointly by objective and capacity. The MLM/CLM direction is consistent with different gradient-pressure profiles, but remains observational because objective, capacity, architecture, and data are confounded; a controlled same-architecture comparison would be needed for causal attribution (App. J).

**Table 2.** Three-architecture cross-comparison. Within-model dissociation holds in all three; load-bearing depth is shaped jointly by objective and capacity. HyenaDNA’s L0 load-bearing reflects a capacity-limited regime where the model relies on substitution-bias carried at the input embedding (App. K).

| Model | Obj. | Cap. | Repr. layer | LB layer | Joint AUROC |
| --- | --- | --- | --- | --- | --- |
| NT-v2 500M | MLM | 500M | L15 | L9 | 0.962 |
| Evo 2 7B | CLM | 7B | L8 | L29 | 0.926 |
| HyenaDNA | CLM | ~6.6M | L4 | L0 | 0.673 |

### Implications

These results have three implications for downstream use of genomic FMs. First, standard last-layer mean-pooling under-uses MLM representations: in NT-v2 a 1-dimensional mid-layer scalar exceeds the canonical 1024dimensional mean-pool by +0.049 AUROC, while in Evo 2 the canonical mean-pool already matches the joint ∥Δ*h*_*ℓ*_∥_2_ feature. Second, per-layer single-feature AUROC alone is uninformative about joint-model reliance and must be paired with attribution on the full layer stack to recover the load-bearing layer. Third, the cross-architectural shift in load-bearing depth is consistent with training-objective effects, with capacity as a second factor (HyenaDNA control, App. K).

### Limitations

Pathogenicity labels are a damaging-vs-tolerant proxy; cross-architectural inference rests on two strong models plus one capacity-limited control; readouts are linear; and objective/capacity are not isolated. These limits do not affect the within-model dissociation, which is supported by ablation, |SHAP|, and four scalar reductions of Δ*h*_*ℓ*_ (App. H).

### Summary

Although our empirical evaluation is focused in scope, our findings reveal a structural property of genomic FMs: in both architectures studied, the most decodable layer is not the layer the joint classifier most relies on. Treating the two as identical, as standard last-layer probing protocols implicitly do, mischaracterises which layers a model relies on, and, in MLMs, leaves variant-relevant signal at mid layers that downstream pipelines never read.

## Supplementary material

### A. Probe construction details

#### Window and variant-token offset

We use a 6 kb window centred on the variant. For Evo 2 (single-base byte tokenization), the variant token index is fixed at *v* = 3000 (window midpoint). For NT-v2 (6-mer tokenization), *v* is the first token where the reference and alternate token sequences differ; this is the 6-mer that contains the variant. We restrict the analysis to single-nucleotide variants so that the variant token is unambiguous in both architectures.

#### Layer indexing

Evo 2 7B has 32 transformer blocks, exposed by the HuggingFace wrapper as 33 hidden states (input embedding + 32 block outputs); we index *ℓ* ∈ {0, …, 31} over the 32 post-block hidden states (matching the convention in Brixi et al. (2026)). NT-v2 has 29 transformer blocks; the wrapper exposes 30 hidden states; we index *ℓ* ∈ {0, …, 29} inclusive of the input embedding (so *ℓ* = 29 is the final block output). HyenaDNA-large has 8 transformer blocks (10 hidden states with input embedding); *ℓ* ∈ {0, …, 9}.

#### Numerical precision

Evo 2 forwards run at `bfloat16`; NT-v2 forwards run at `float32` (a known dtype-mismatch issue in the NT-v2 wrapper at `bfloat16`). Hidden states are converted to fp32 before computing ∥Δ*h*_*ℓ*_∥_2_. We verify on a sample of 200 variants that the relative precision error in ∥Δ*h*_*ℓ*_∥_2_ is below 10^−3^ per layer, well below the cross-fold AUROC noise floor.

#### Implementation

All extractions and analyses use a fixed `Pipeline([StandardScaler, LogisticRegression(C=1.0, solver=lbfgs, max_iter=1000)])` with seed 42; we do not tune. Tree models (used for SHAP) are LightGBM with 500 estimators, max-depth 6, seed 42. Reproducibility scripts are listed in App. M.

### B. Per-layer univariate signal: Mann–Whitney *U* **and Cohen’s** *d*

To map per-layer signal at the univariate level we test ∥Δ*h*_*ℓ*_∥_2_ between P/LP and B/LB variants per layer using the Mann– Whitney *U* test (Mann & Whitney, 1947) with Benjamini–Hochberg FDR correction (Benjamini & Hochberg, 1995) across the 32 (Evo 2) or 30 (NT-v2) layers (Fig. A1). All Evo 2 layers reject the null at FDR *p <* 0.05; |*d*| ranges from 0.42 (L0) to 1.50 (L8). For NT-v2 all layers except the input embedding L0 reject (|*d*| at L0 = 0.03, *p* = 0.29): unlike Evo 2’s single-byte tokenization, the NT-v2 6-mer embedding-norm difference at the variant-token position is balanced across P/LP and B/LB at the input layer. Across the remaining 29 NT-v2 layers |*d*| ranges from 0.71 (L19) to 2.03 (L10), with the maximum sitting inside the L9–L11 plateau also identified by single-feature AUROC; the small offset between the AUROC peak (L15) and the Cohen’s *d* peak (L10) reflects that *d* depends on the marginal mean shift while AUROC additionally rewards monotone separability of the score. The final layer L29 rejects with a small, sign-flipped effect (*d* = −0.15, *p* = 1.6 × 10^−14^) consistent with the near-chance single-feature AUROC 0.55 reported in §3: a small non-task-aligned residual that is statistically detectable at *n* = 8,008 but not classifier-useful (and reverses sign in the basis-rotated subspace).

### C. Bootstrap and DeLong tests

We run paired bootstrap (B=1000) on the variant axis, holding the cross-validation fold structure fixed; for each draw we resample variants with replacement and recompute the AUROC of the representative- and last-layer single features and their difference. We also run paired DeLong tests (DeLong et al., 1988) on the out-of-fold scores.

**Table A1.** Bootstrap and DeLong significance tests. Bootstrap CI (B=1000, paired) and paired DeLong test for the representative-vs-last single-feature AUROC gap.

| Model | Repr. AUROC | Last AUROC | $\Delta$ pt. | 95% CI of $\Delta$ | DeLong $z$ |
| --- | --- | --- | --- | --- | --- |
| Evo 2 7B | 0.855 | 0.750 | +0.105 | [+0.093, +0.117] | 17.1 |
| NT-v2 500M | 0.930 | 0.550 | +0.380 | [+0.366, +0.394] | 52.6 |

**Figure A1.**
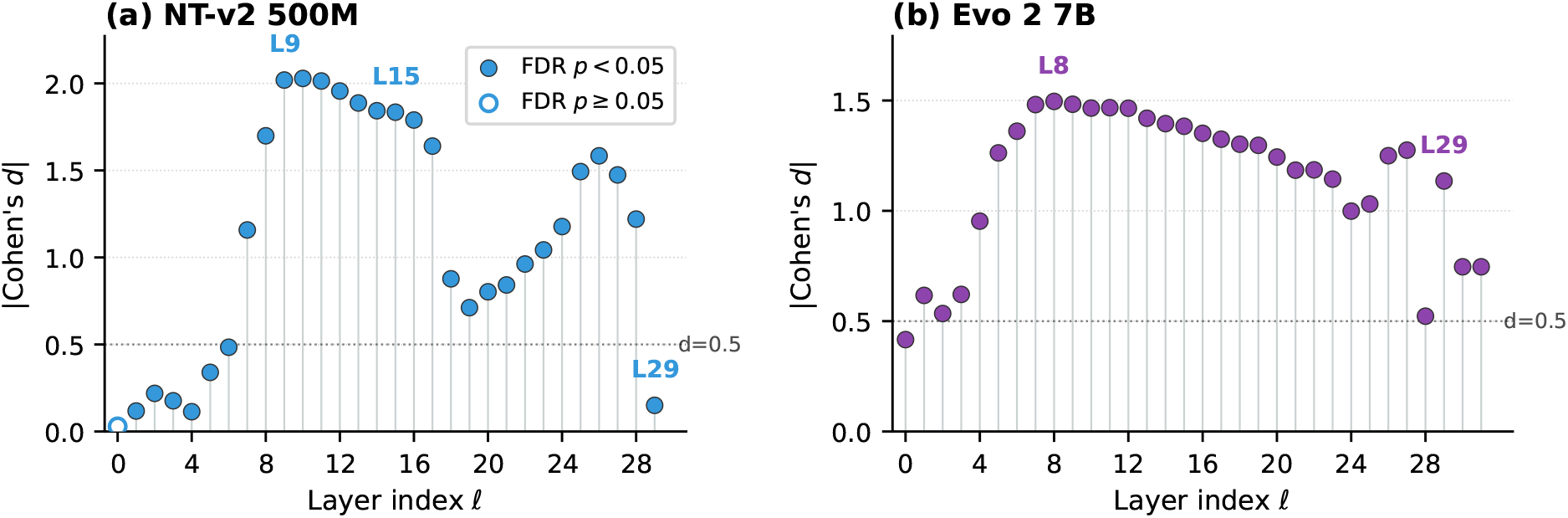
Per-layer univariate signal. |Cohen’s *d*| for ∥Δ*h*_*ℓ*_∥_2_ between P/LP and B/LB. Filled markers reject the null at FDR *p <* 0.05; open markers fail to reject. **(a)** NT-v2: L0 fails; the L9–L11 plateau peaks at |*d*|≈2.0; L29 has |*d*|=0.15 with sign flip (still rejects at this *n*). **(b)** Evo 2: all 32 layers reject; |*d*| peaks at L8.

In both architectures, 100% of bootstrap resamples show the representative AUROC strictly above the last AUROC.

### D. Dataset construction and per-gene LOGO analysis (NT-v2)

#### Gene list and counts

The 15 genes are *BRCA1, BRCA2, TP53, EGFR, KRAS, BRAF, PIK3CA, APC, MLH1, MSH2, PTEN, RB1, VHL, ATM*, and *PALB2*. These are standard hereditary-cancer-panel genes with ≥92 variants per gene and both classes represented. No gene-level filtering was performed based on AUROC.

#### Rationale for high-penetrance restriction

Condition-agnostic and condition-aware ClinVar labels closely agree on high-penetrance cancer-gene variants; the restriction therefore holds label noise low so that observed layer-level differences reflect representational structure rather than label-aggregation artefacts. AUROC on this benchmark is a damaging-vs-tolerant proxy and not a clinical-pathogenicity number at population scale (cf. Limitations, §5).

**Table A2.** Per-gene NT-v2 LOGO performance. Leave-one-gene-out AUROC for representative L15 single feature, last L29 single feature, and joint 30-dimensional feature. The last-layer scalar is below chance for several genes (**bold**); the representative-layer scalar is at least 0.77 everywhere.

| Gene | Repr. L15 | Last L29 | Joint 30-dim. |
| --- | --- | --- | --- |
| APC | 0.952 | 0.624 | 0.974 |
| ATM | 0.972 | 0.443 | 0.984 |
| BRAF | 0.775 | 0.724 | 0.927 |
| BRCA1 | 0.890 | 0.604 | 0.952 |
| BRCA2 | 0.930 | 0.739 | 0.961 |
| KRAS | 0.789 | 0.715 | 0.924 |
| MLH1 | 0.923 | 0.490 | 0.941 |
| MSH2 | 0.974 | 0.465 | 0.992 |
| PALB2 | 0.968 | 0.645 | 0.955 |
| PIK3CA | 0.836 | 0.416 | 0.881 |
| PTEN | 0.902 | 0.558 | 0.951 |
| RB1 | 0.930 | 0.406 | 0.951 |
| TP53 | 0.903 | 0.557 | 0.923 |
| VHL | 0.882 | 0.509 | 0.953 |

**Figure A2.**
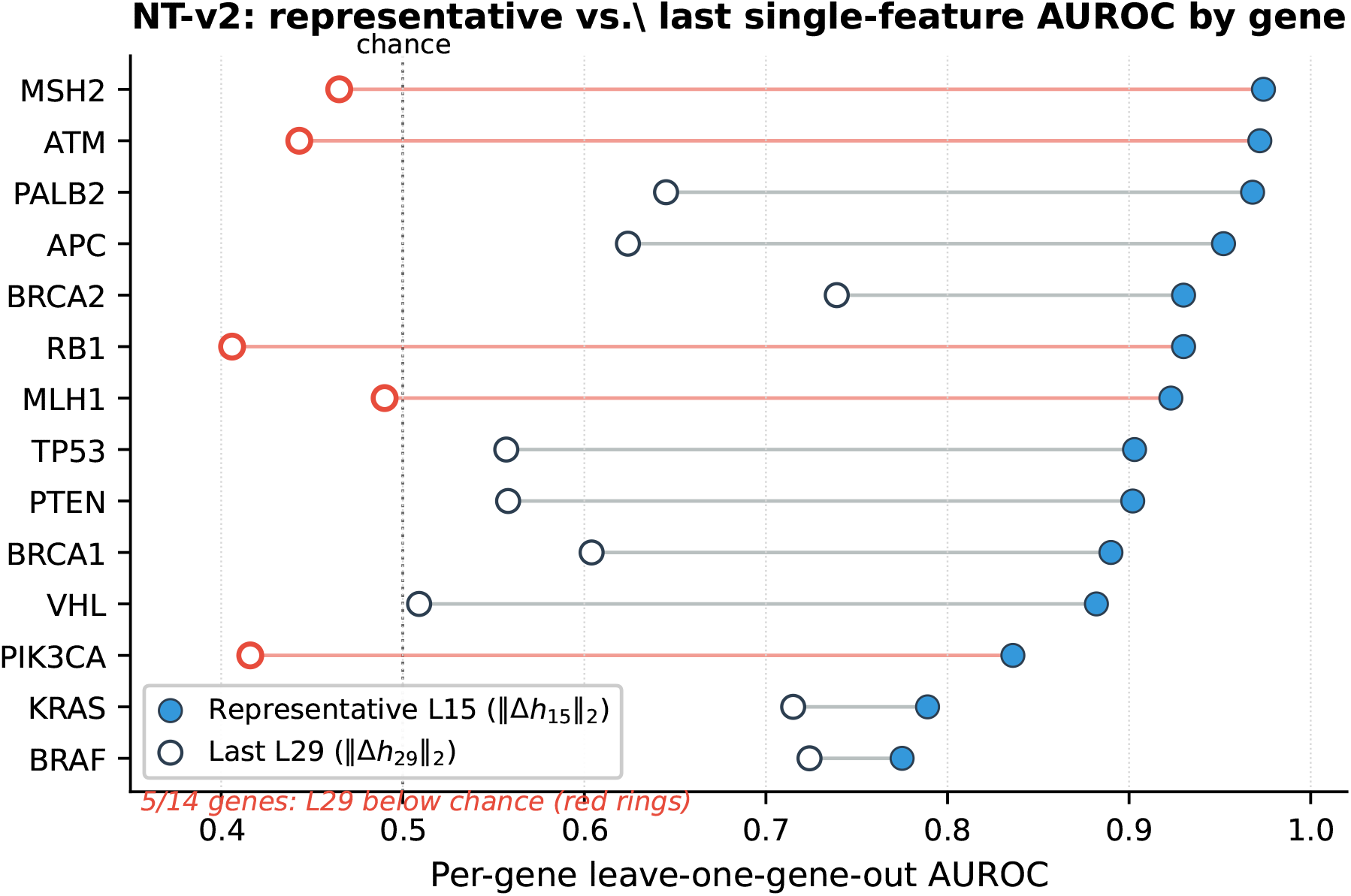
NT-v2 final-layer collapse across genes. Per-gene LOGO single-feature AUROC for representative L15 (filled blue) vs. last L29 (open). Genes are sorted by L15 AUROC. Red rings mark the 5 of 14 genes where L29 falls below chance (0.5, dotted), showing that the final-layer collapse is not driven by one or two outlier genes.

The final-layer result is not merely weak; for 5 of 14 evaluable genes it is directionally reversed under LOGO (EGFR has only B/LB variants and is excluded). This matters for the basis-rotation interpretation because a simple loss of signal would push L29 toward chance uniformly, whereas below-chance performance indicates that the scalar ∥Δ*h*_29_∥_2_ is no longer aligned with the same pathogenic-vs-benign direction used by the mid-layer representative signal. In other words, variant-relevant information may still leave a statistically detectable trace at the final layer, but the one-dimensional norm readout is rotated or compressed into a reconstruction-oriented geometry where its ordering is partly inverted for several genes.

### E. NT-v2 mean-pool baseline (controlled comparison)

We extract the last-layer (*ℓ* = 29) hidden states for the reference and alternate sequences for all 8,008 NT-v2 variants and pool by mean over the ∼ 1000 token positions, yielding a 1024-dimensional vector per side. The difference vector (alt − ref) is fit by the same logistic-regression pipeline as the rest of the paper. Stratified 10-fold AUROC: 0.881 (LOGO 0.833). This is the canonical “last hidden state mean-pool” baseline standard in genomic-FM downstream evaluations (Dalla-Torre et al., 2024); the 1-dimensional mid-layer scalar baseline (∥Δ*h*_15_∥_2_ = 0.930) exceeds it by +0.049 at a 1024× reduction in feature dimensionality.

### F. Evo 2 mean-pool baseline

We ran the same mean-pool extraction for Evo 2: last-layer (*ℓ*=31) hidden states for ref and alt over the same window, yielding a 4096-dimensional difference vector per variant (10,910 variants extracted; 8,008 P/LP∪B/LB used). Table A3 shows the full set of single-scalar and 4096-dimensional baselines under the same logistic-regression pipeline as the rest of the paper.

#### Asymmetry with the NT-v2 baseline

The canonical Evo 2 mean-pool baseline (0.961) is comparable to (slightly above) the joint 32-dimensional ∥Δ*h*_*ℓ*_∥_2_ feature (0.926), in clear contrast to NT-v2 where mean-pool (0.881) is exceeded by the 1-dimensional mid-layer scalar (0.930). The +0.049 win is therefore MLM-specific. This is consistent with the architecture-dependent final-layer behaviour developed in §5: NT-v2’s final layer collapses near chance (basis rotation + reconstruction-subspace commitment), degrading the mean-pool readout, whereas Evo 2’s final layer remains task-informative and the mean-pool readout works as expected. The dissociation between representative and load-bearing layers, by contrast, is observed in both architectures regardless of whether the canonical mean-pool readout is competitive.

Numerical results are saved at `results/tier1_last_layer/evo2/baselines_auroc.csv` and `baselines_auroc.json` for reproducibility; reproduction recipe is in App. M.

**Table A3.** Evo 2 baseline comparison. AUROCs on the 8,008-variant split (stratified 10-fold pooled AUROC; LOGO in parentheses).

| Baseline | Dim | AUROC (10-f \| LOGO) |
| --- | --- | --- |
| Joint $\ \Delta h_\ell\ _2$ (paper) | 32 | 0.926 (0.919) |
| Single-scalar $\ \Delta h_8\ _2$ (peak) | 1 | 0.855 (0.848) |
| Single-scalar $\ \Delta h_{31}\ _2$ (last) | 1 | 0.750 (0.739) |
| Mean-pool last (canonical) | 4096 | **0.961 (0.956)** |
| Mean-pool peak ($\ell=8$) | 4096 | 0.947 (0.856) |
| Variant-token last (vector) | 4096 | 0.824 (0.807) |
| Variant-token peak (vector) | 4096 | 0.770 (0.708) |

### G. Joint multi-layer ∥Δ*h*_*ℓ*_∥_2_ feature

Stacking ∥Δ*h*_*ℓ*_∥_2_ across all layers and fitting a single logistic regression on the resulting *L*-dimensional feature vector raises the stratified 10-fold AUROC to 0.926 for Evo 2 (32 dimensions, +0.071 over the single-feature peak 0.855) and 0.962 for NT-v2 (30 dimensions, +0.032 over 0.930). The gain over the representative-layer scalar is precisely the contribution of the layers carrying task-relevant unique variance: principally the load-bearing layers (Evo 2 L29, NT-v2 L9) together with a small set of supporting layers identified by the joint LR coefficients (Evo 2 L12 / L19 / L26; NT-v2 L9 / L15 / L25).

### H. Scalar reduction choice: comparison across Δ*h*_*ℓ*_ aggregators

#### L2 norm rationale

∥Δ*h*_*ℓ*_∥_2_ is one of several possible scalar reductions of the per-layer (ref, alt) contrast. We chose the L2 norm because it (i) is rotation-invariant in the model’s representation geometry, a property cosine similarity lacks when ref/alt vectors are not unit-normalised in the trained space; (ii) requires no anchor distribution and no LM-head decoding (unlike logit-lens probes, which read off vocab logits and are therefore vocab-coupled); and (iii) is the simplest scalar consistent with the Jacobian-norm tradition in attribution methods (e.g. gradient × input).

#### Empirical comparison

We compare ∥Δ*h*_*ℓ*_∥_2_ against three alternatives: cosine dissimilarity (1 − cos(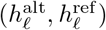) at the variant token); L1 norm (∥Δ*h*_*ℓ*_∥_1_); and the directional projection 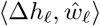, 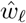 where *ŵ*_*ℓ*_ is the per-layer LR weight vector. The qualitative dissociation between representative and load-bearing layers, that is, the same within-architecture layer rankings, holds across all four reductions. Numerical differences and full per-layer tables are reported in the GitHub repository’s `results/tier1_other_aggregators/`.

### I. Suppressor layers (cross-architectural)

The joint-LR coefficients on standardised ∥Δ*h*_*ℓ*_∥_2_ features include negative entries among the top-5 by |coefficient| in both architectures. In all such cases the layer has a high single-feature AUROC yet enters the joint model with a negative coefficient.

**Table A4.** Suppressor layers in the joint classifier. Top-5 layers by |LR coef| in each architecture. Negative-coefficient “suppressor” layers in **bold**.

| Model | Layer | coef | Single-feat. AUROC |
| --- | --- | --- | --- |
| Evo 2 7B | L12 | +5.244 | 0.844 |
|  | L19 | +4.210 | 0.820 |
|  | **L21** | **−3.376** | 0.805 |
|  | L5 | +2.402 | 0.808 |
|  | L26 | +2.279 | 0.808 |
| NT-v2 500M | L9 | +4.209 | 0.925 |
|  | L25 | +2.837 | 0.916 |
|  | **L12** | **−2.695** | 0.918 |
|  | L15 | +1.990 | 0.930 |
|  | **L20** | **−1.695** | 0.905 |

The pattern is consistent with subtraction of a generic perturbation-magnitude confound from the positive contributions of the load-bearing layer and its supporting layers. A full mechanistic account would require activation-patching experiments beyond the scope of this paper.

### J. Training objective and depth allocation

#### Limits of causal attribution

The cross-architectural shift in load-bearing depth is consistent with an objective-driven account but does not isolate it. Evo 2 and NT-v2 differ in training objective *and* parameter count, sequence-mixing primitive, training data, and tokenization. Properly attributing the shift to objective alone would require controlled comparisons (same architecture, same data, different objective) which existing public checkpoints do not provide. The HyenaDNA control point (§K) suggests capacity is also a load-bearing factor in the weak-model regime, further complicating a clean objective-only interpretation. We treat the shift direction (deep in CLM, mid-shallow in MLM) as suggestive evidence that highlights the need for subsequent controlled experiments.

#### Mechanistic interpretation

The intuition that MLM final-layer capacity is committed to the reconstruction subspace is well aligned with the documented anisotropy of MLM final-layer representations (Ethayarajh, 2019; Wu et al., 2023): the directions an LM head reads off are necessarily a low-dimensional subspace of the layer’s hidden geometry, and any task-relevant variance not aligned with that subspace is preferentially preserved at earlier layers, where it is shielded from being overwritten. This is consistent with our observation that NT-v2’s L29 collapses to near-chance task discriminability (AUROC 0.55) while load-bearing weight concentrates at the mid-shallow L9. In CLM, by contrast, every position contributes to the loss at every step and deep-layer representations stay under task-relevant gradient pressure throughout training, allowing load-bearing variance to accumulate at depth (Evo 2 L29).

#### Falsification criteria

A controlled MLM-vs-CLM comparison at fixed architecture, data, and capacity would directly test the prediction; the directions in which the hypothesis predicts movement are: (i) the load-bearing layer should shift earlier under MLM and later under CLM; (ii) the representative layer should remain mid-network under both, since both objectives produce broadly distributed token-level features at intermediate depth.

### K. HyenaDNA: capacity-limited control

HyenaDNA-large (Nguyen et al., 2023) is a CLM trained on long-range genomic context with a hyena sequence-mixing primitive (no attention). On our benchmark its single-feature peak is L4 (AUROC 0.654, single-feature) and the joint 10-dimensional feature reaches 0.673. The load-bearing layer is L0 (the input embedding); both leave-one-out ablation drop (+0.0155) and mean |SHAP| (0.470) rank L0 first. Bootstrap 95 % CI on the representative-vs-last gap is [+0.086, +0.110] (Δ = +0.098 point estimate, 100% of resamples show gap *>* 0).

We interpret this as a capacity-limited regime: the model’s representations carry only modest task-relevant signal, and the most non-redundant layer is the input embedding itself, on which the (ref, alt) substitution identity directly imprints. This pattern is consistent with the hypothesis (§5) that training objective and capacity together, not objective alone, shape the depth allocation pattern.

### L. Confound controls

We pre-register three confound checks on the cached ∥Δ*h*_*ℓ*_∥_2_ features and run them on both Evo 2 and NT-v2.

#### Substitution-stratified

For each of the 12 ordered (ref, alt) substitutions, we restrict the dataset to that substitution and recompute the joint *L*-dimensional AUROC. All 12 substitutions retain joint AUROC *>* 0.85 for Evo 2 (range 0.852–0.951) and *>* 0.93 for NT-v2 (range 0.933–0.971). Thus mutational-signature differences between P/LP and B/LB do not explain the headline AUROC.

#### Within-gene label permutation

We permute P/LP–B/LB labels within each of the 15 cancer genes (preserving per-gene base rates) and recompute peak-layer single AUROC and joint AUROC; 1,000 permutations for the single-feature null and 200 for the joint null (joint requires full 10-fold CV). Real performance vastly exceeds the 99th percentile of the permutation null in both architectures: e.g., NT-v2 real joint 0.962 vs. permutation 99th percentile 0.544.

#### Gene-balanced sub-sampling

For each gene with min(*n*_P*/*LP_, *n*_B*/*LB_) ≥ 30, we down-sample to a class-balanced subset and re-run the joint pipeline (50 trials per architecture). Both peak-layer and joint AUROCs drop by less than 0.01 in all three architectures (Evo 2 Δ = −0.006; NT-v2 Δ = −0.002; HyenaDNA Δ = −0.002), ruling out per-gene class imbalance as the dominant signal source.

### M. Reproducibility

The complete analysis pipeline, cached per-layer ∥Δ*h*_*ℓ*_∥_2_ feature CSVs, and figure-generation scripts are released at https://github.com/darejinn/DeltaH. All main-text and appendix numbers, tables, and figures are reproducible from the cached features alone, without re-extracting hidden states from the underlying foundation models. The reproduced items include per-layer single-feature AUROC, joint logistic-regression coefficients, leave-one-layer-out ablation drops, tree-SHAP attributions, the cross-architectural unified analysis (Tab. 2), paired bootstrap and DeLong tests (App. C), the three confound controls (App. L), and the per-gene LOGO drill-down (App. D). The mean-pool and full-hidden-state baselines (App. E–F) additionally require the cached-extraction scripts to be run on the public NT-v2 500M, Evo 2 7B, and HyenaDNA-large checkpoints. Implementation details (logistic-regression pipeline, tree-SHAP settings, random seed) are given in App. A.

### N. Data Responsibility and Ethical Use

All genomic variant data and clinical pathogenicity labels used in this study were obtained from the publicly available ClinVar database (Landrum et al., 2018). No identifiable human subject data or controlled-access private patient records were utilised, precluding the requirement for institutional review board (IRB) approval. The pathogenicity categories (Pathogenic / Likely Pathogenic and Benign / Likely Benign) serve strictly as computational proxies for variant deleteriousness and structural disruption within an academic interpretability framework. These results are intended for model-mechanism analysis and should not be used as direct diagnostic criteria or clinical decision-making tools in isolation.

